# Distal Protein-Protein Interactions Contribute to SARS-CoV-2 Main Protease Substrate Binding and Nirmatrelvir Resistance

**DOI:** 10.1101/2024.04.01.587566

**Authors:** Eric M. Lewandowski, Xiujun Zhang, Haozhou Tan, Aiden Jaskolka-Brown, Navita Kohaal, Aliaksandra Frazier, Jesper J. Madsen, Lian M.C. Jacobs, Jun Wang, Yu Chen

## Abstract

SARS-CoV-2 main protease, M^pro^, is responsible for the processing of the viral polyproteins into individual proteins, including the protease itself. M^pro^ is a key target of anti-COVID-19 therapeutics such as nirmatrelvir (the active component of Paxlovid). Resistance mutants identified clinically and in viral passage assays contain a combination of active site mutations (e.g. E166V, E166A, L167F), which reduce inhibitor binding and enzymatic activity, and non-active site mutations (e.g. P252L, T21I, L50F), which restore the fitness of viral replication. Although the mechanism of resistance for the active site mutations is apparent, the role of the non-active site mutations in fitness rescue remains elusive. In this study, we use the model system of a M^pro^ triple mutant (L50F/E166A/L167F) that confers not only nirmatrelvir drug resistance but also a similar fitness of replication compared to the wild-type both in vitro and in vivo. By comparing peptide and full-length M^pro^ protein as substrates, we demonstrate that the binding of M^pro^ substrate involves more than residues in the active site. In particular, L50F and other non-active site mutations can enhance the M^pro^ dimer-dimer interactions and help place the nsp5-6 substrate at the enzyme catalytic center. The structural and enzymatic activity data of M^pro^ L50F, L50F/E166A/L167F, and others underscore the importance of considering the whole substrate protein in studying M^pro^ and substrate interactions, and offers important insights into M^pro^ function, resistance development, and inhibitor design.

## Main

Since its emergence in late 2019, the COVID-19 pandemic, caused by severe acute respiratory syndrome coronavirus 2 (SARS-CoV-2), has had immense health, economic, and social impacts globally. With new infection rates remaining high^1^, SARS-CoV-2 continues to pose an urgent health threat. The main protease (M^pro^) of SARS-CoV-2 is a homo-dimeric cysteine protease responsible for cleaving the viral polyproteins into its mature substituents, including M^pro^ itself, during viral replication^2,3^. M^pro^ is a validated drug target and is inhibited by nirmatrelvir (PF-07321332)^4^, which is the active agent in Pfizer’s oral COVID-19 drug, Paxlovid (nirmatrelvir/ritonavir combination)^5^. While Paxlovid is one of the most effective COVID-19 treatments to date^6^, M^pro^ has demonstrated that it is prone to mutations in, and outside, the active site, which has resulted in the rise of SARS-CoV-2 variants showing resistance to nirmatrelvir^7,8^.

Recent studies have shown that the SARS-CoV-2 virus harboring a M^pro^ triple mutant, L50F/E166A/L167F, is highly resistant to nirmatrelvir while demonstrating similar fitness of replication as the wild-type (WT) virus in cell culture and animal models^9–11^. L50F locates away from the M^pro^ active site and is not directly involved with binding to nirmatrelvir or the nsp4-5 peptide substrate commonly used in biochemical analysis^8,12,13^. Yet, L50F is able to rescue the reduced viral fitness caused by the active site substitutions (e.g., E166A, E166V)^14^. Biochemical analysis has indicated that compensatory mutations outside the active site, such as L50F, have little impact on the hydrolysis of the viral nsp4-5 peptide substrate, and improve the enzymatic activity of the active site mutants, such as E166V, by only two-fold (from 3% of WT activity for E166V to 6% for L50F/E166V), yet they fully restore the viral fitness in cell-based studies^15^. The discrepancy between the biochemical data and viral replication assay remains a puzzle and impedes our ability to understand M^pro^ resistance mutations and to develop new inhibitors.

The majority of the in vitro biochemical studies to date have only used small peptide substrates of SARS-CoV-2 corresponding to the nsp4-5 M^pro^ cleavage site on the viral polyproteins, which corresponds to the M^pro^ N-terminal self-cleavage site. Whereas the peptides can capture the enzyme-substrate (ES) interactions inside the active site, we hypothesize that there may be important protein-protein interactions outside the active site in the ES complex. By only using small peptide substrates, and not the whole protein, critical protein-protein interactions during ES formation may be missed that can help to explain how these distal mutations assert their effects. Here, we show that a more holistic approach may be needed when studying the M^pro^ enzymatic function and the effects of resistance mutations.

In order to determine the effect of the different M^pro^ mutations on substrate binding and nirmatrelvir resistance, we characterized the L50F/E166A/L167F triple mutant using an enzymatic assay and X-ray crystallography. Using the nsp5-6 cleavage sequence containing FRET peptide and the full-length M^pro^ as substrates, we determined the catalytic activity of the L50F/E166A/L167F triple mutant and compared the results with the WT, as well as corresponding single and double mutants. Additionally, the crystal structures of M^pro^ L50F/E166A/L167F and M^pro^ L50F were determined to 2.23 and 2.21 Å (Supplementary Data Table 1), respectively, to illustrate the underlying molecular interactions.

### Effects of distal mutations on M^pro^ activity using protein substrates

M^pro^ cleaves the viral polyproteins at 11 sites, including those at its own two termini. We posit that the rate-limiting step in the processing of the viral polyproteins by M^pro^ is the self-cleavage of the protease from the viral polyprotein, namely the M^pro^ N-terminal nsp4-5 sequence, and the C-terminal nsp5-6 sequence (Figure 1a). A recent study has shown that M^pro^ self-cleavage is intra-molecular at its N-terminus (nsp4-5), and inter-molecular at its C-terminus (nsp5-6), involving an M^pro^ dimer functioning as enzyme acting on another M^pro^ dimer serving as substrate (Figure 1b)^16^. This suggests that the cleavage at its C-terminus (nsp5-6) may be more susceptible to inhibitors such as nirmatrelvir. Hence, the most effective resistance mutations may enhance the enzymatic activity at this site. It is consistent with the observation that a common resistance mutation, T304I, occurs at the M^pro^ C-terminus cleavage site. However, the mechanism of T304I is not entirely clear. Moreover, the enzymatic activity of M^pro^ in digesting the nsp5-6 sequence has not been characterized for any of the resistance mutants.

**Figure 1.**
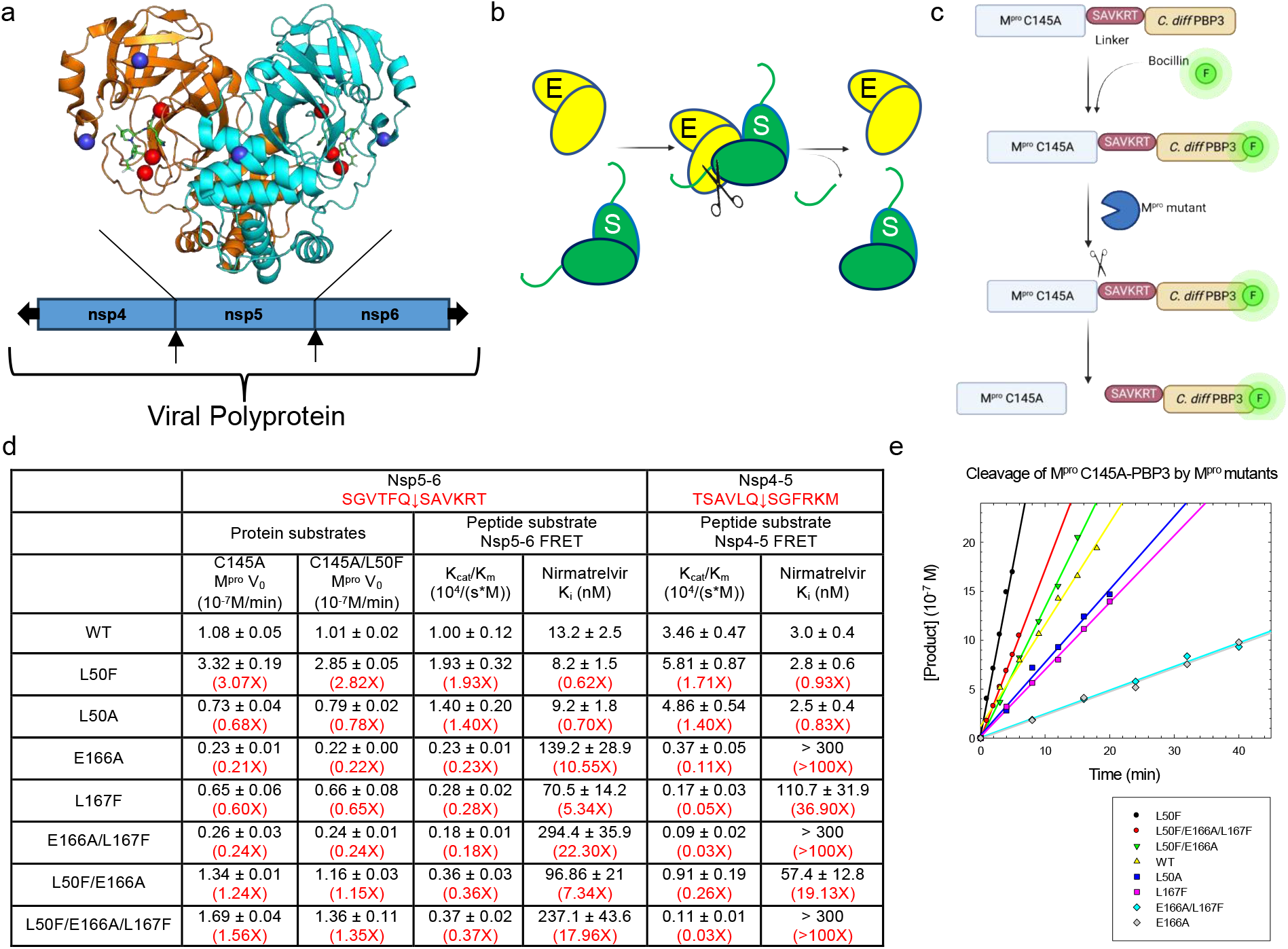
Characterization of M^pro^ activity using nsp5-6 substrates. **a)** Overview of the M^pro^ dimer showing nirmatrelvir (green) bound in the active site and the locations of mutations of interest in the active site (red spheres, e.g., E166/L167) and distal to the active site (blue spheres, e.g., L50) (PDB 8DCZ). **b)** M^pro^ self-cleavage at the C-terminus. E: M^pro^ as enzyme, S: M^pro^ as substrate. The polyprotein chain after M^pro^ is represented by the curved line. **c)** Schematic of how the fluorescence gel-based assay functions. **d)** Rate of activity for the C145A and the C145A/L50F mutants. **e)** Cleavage reaction of C145A-PBP3 by multiple M^pro^ mutants. The cleavage reaction of C145A/L50F-PBP3 by the same M^pro^ mutants is shown in Supplementary Data Fig. 1.

To determine the effect that the nirmatrelvir-resistant triple mutant L50F/E166A/L167F had on M^pro^ activity, we employed a novel fluorescence electrophoresis-based assay using a M^pro^ protein substrate in which the catalytically inactive M^pro^ C145A protein was linked to *Clostridium difficile* Penicillin Binding Protein 3 (PBP3) by the first six residues of the nsp6 N-terminus (Figure 1c), which, together with the M^pro^ C-terminal residues, corresponds to the nsp5-6 cleavage sequence. The majority of the nsp6 protein is membrane-embedded and thus not included in the fusion construct. A fluorescently labeled penicillin, Bocillin, was used to covalently react with the catalytic serine of PBP3 and monitor the ability of M^pro^ enzyme to cleave at the nsp5-6 junction of the fusion protein. By monitoring the fluorescence intensity of the band representing the “cut” nsp6 linked PBP3 on a gel, we were able to determine the activity of the M^pro^ mutants (Figure 1d-e, Supplementary Data Figure 1-3). The concentrations for both the M^pro^ enzyme and substrate M^pro^ fusion protein were fixed in our experiments, and the initial velocity of the reaction was determined and compared. At high substrate concentrations, interactions among the fusion protein substrate molecules appeared to sequester the cleavage site and result in a unique ‘substrate inhibition’. Therefore, we were unable to vary the substrate concentrations and obtain *k_cat_/K_m_* values of the reaction. In parallel to the protein substrate assay, we also characterized the enzymatic activity and nirmatrelvir inhibition of the M^pro^ L50F/E166A/L167F triple mutant using the conventional FRET substrates containing the 12-residue nsp5-6 or nsp4-5 cleavage sequence.

In our experiments, we focused on the L50F/E166A/L167F triple mutant and the corresponding single and double mutants (e.g., E166A/L167F), as this triple mutant is one of the most clinically relevant nirmatrelvir drug-resistant mutants with a similar fitness of replication as the WT. Compared with the experiments using the peptide substrates (nsp4-5 and nsp5-6 FRET peptides), L50F exhibited significantly larger effects on the M^pro^ enzymatic activity when protein substrates were used (M^pro^ C145A and C145A/L50F fusion proteins) (Figure 1d). In our experiments, M^pro^ L50F was found to be ∼3 times more active than the WT in cleaving the C145A M^pro^ protein substrate, while it was 1.93- and 1.71-fold more active than the WT in cleaving the nsp5-6 and nsp4-5 FRET substrates, respectively. In comparison, the L50A mutant was less active in cleaving the C145A and C145A/L50F M^pro^ protein substrates (0.68-fold), while showing slightly more activity than the WT (1.40-fold) in the peptide substrate assays. The E166A and L167F single mutants, and the E166A/L167F double mutant, all had reduced enzymatic activity in cleaving the nsp5-6 peptide and protein substrates (0.18 to 0.65-fold of WT). In comparison, a more profound reduction of enzymatic activity was observed in cleaving the nsp4-5 FRET peptide substrate (0.03 to 0.1-fold of WT), suggesting the nsp5-6 cleavage sequence is more relevant in explaining the restoration of the fitness of replication. This difference between the peptide substrates appears to have largely originated from the L167F mutation, which showed 0.28- and 0.05-fold activity of the WT for the nsp5-6 and nsp4-5 peptides, respectively. The most striking observation from our studies was the ability of L50F (3.07-fold) to rescue the reduced enzymatic activity of the E166A/L167F double mutant (0.24-fold) with the L50F/E166A/L167F triple mutant (1.56-fold) exhibiting slightly better activity than WT in cleaving the C145A M^pro^ protein substrate. In comparison, L50F was less effective in rescuing the reduced enzymatic activity of the E166A/L167F double mutant in cleaving the nsp5-6 (0.18-fold to 0.37-fold WT) and nsp4-5 (0.03- fold to 0.03-fold) FRET peptide substrates. Since the L50F mutation may affect protein-protein interactions when present on either the enzyme or protein substrate, we also investigated the reactions using C145A/L50F mutant in the fusion substrate protein. Overall, the results were similar to that of the C145A substrate, suggesting that L50F exerts its influence on the reaction mainly on the protein substrate rather than the peptide substrate. Collectively, our results showed that the L50F mutant can rescue the reduced enzymatic activity of the E166A/L167F mutant to a similar level as the WT protein when the M^pro^ protein with the nsp5-6 cleavage sequence was used as the substrate. Compared to the low activity of the triple mutant in hydrolyzing the nsp4-5 peptide substrate assay (3% of WT), our results using the M^pro^ protein substrates highlight the differences between the two assays and the importance of using protein substrates in studying M^pro^ resistance mutations.

### L50F/E166A/L167F triple mutant crystal structure

The M^pro^ triple mutant crystallized in the P2_1_ space group at 2.23 Å resolution, with four copies of the protein in the asymmetric unit, with each biological dimer forming a “dimer of dimers” together with another M^pro^ dimer from an adjacent asymmetric unit (Figure 2a). Interestingly, the side chain of the mutated F50 residue of one M^pro^ protomer resides at this dimer-dimer interface and nestles into a hydrophobic pocket that is formed by V212, L220, T257, and I259 residing on helices of an adjacent M^pro^ protomer from a different dimer. Moreover, the C-terminus of this M^pro^ protomer from the adjacent dimer enters the active site inside the M^pro^ protein harboring the aforementioned F50 from the bottom of the catalytic pocket, similar to what is observed in the complex structure of M^pro^ with peptide substrates^17^. These observations suggest that the triple mutant crystal structure has captured the product complex of M^pro^ self-cleavage and that the L50F mutation may promote protein-protein interactions facilitating the positioning of the C-terminal substrate peptide in the active site for cleavage. It is also possible that the aromatic fluorophore of the FRET peptide substrate may mimic some of these inter-molecular interactions with F50, leading to enhanced substrate binding and increased reaction rate.

**Figure 2.**
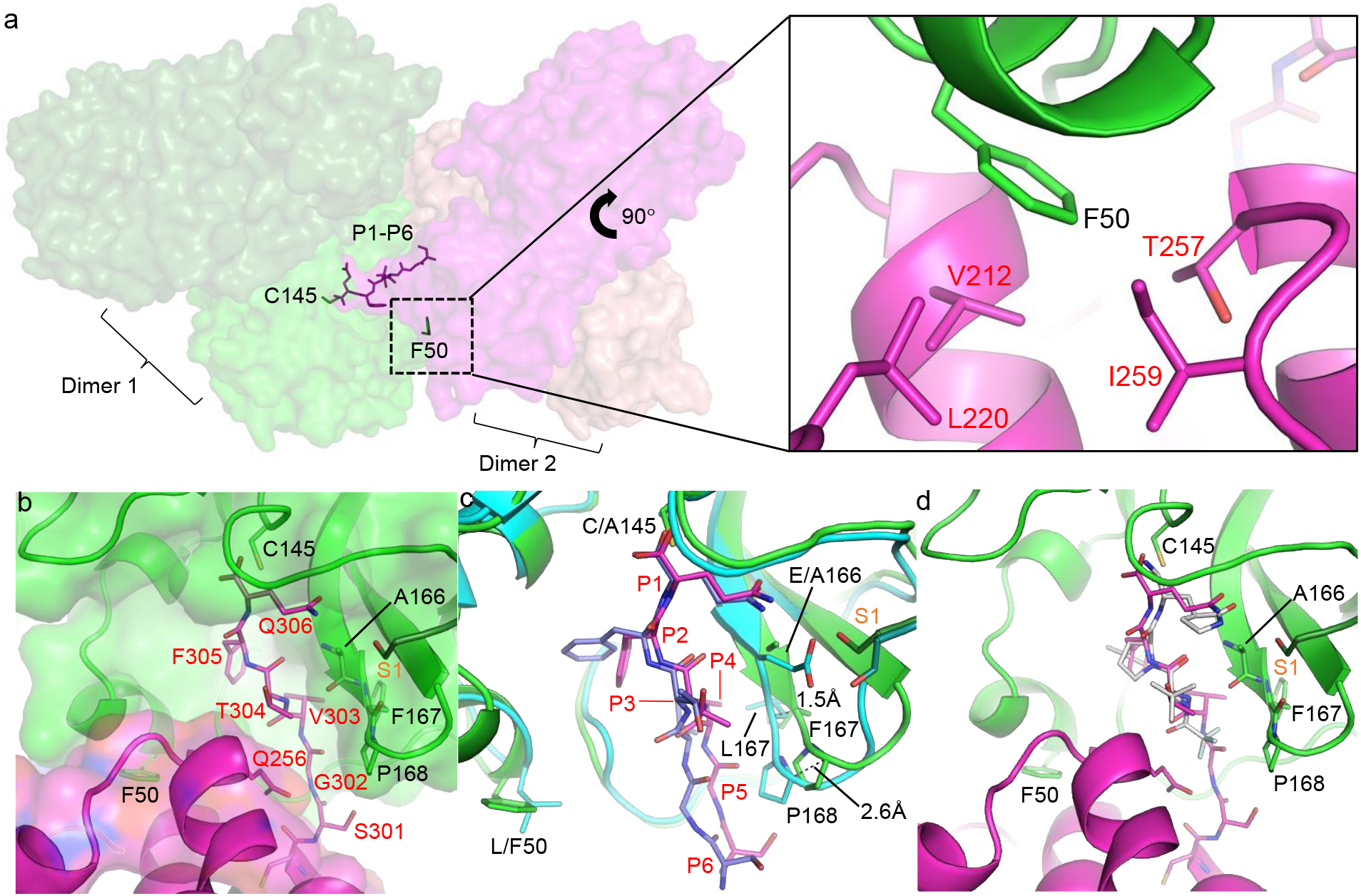
M^pro^ L50F/E166A/L167F triple mutant. **a)** M^pro^ triple mutant dimer of a dimer (dark green/green, magenta/salmon), showing P1-P6 (magenta stick) of the C-terminus from the substrate dimer bound in the active site of the enzyme dimer (green). Zoomed in view shows the interactions of the enzyme F50 (green) within the substrate hydrophobic pocket (magenta). Substrate residues are noted in red text. **b)** Close up of the binding pose of P1-P6 bound in the triple mutant active site (substrate shown in magenta and enzyme shown in green). The N-terminus from an adjacent monomer is noted in orange. Substrate residues are noted in red text. **c)** Movement of the 166-168 backbone in the triple mutant with P1-P6 bound in the active site (magenta/green) vs. M^pro^ C145A with nsp5/6 substrate bound (cyan/light purple) (PDB 7MB5). Substrate residues are noted in red text, and the N-terminus from an adjacent monomer is noted in orange. **d)** Binding pose of nirmatrelvir (white, PBD 8DCZ) superimposed into the triple mutant binding pocket. The N-terminus from an adjacent monomer is noted in orange.

In addition to the intermolecular interactions, intramolecular interactions between the C-terminus and the Q256-containing helix of the same protomer are also observed and help the placement of the substrate peptide. Specifically, these intramolecular interactions create a new well-defined S3 site to accommodate T304, which forms contacts with the helix backbone surrounding Q256 (Figure 2b-d). These interactions were not present in previous complex structures with various peptide substrates, and they help explain the resistance mutation of T304I, as an isoleucine side chain can enhance the non-polar contacts between residue 304 and the Q256 side chain and backbone atoms. Interestingly, Q256L was also observed in the viral passage assays^15^, and can enhance the intramolecular interactions with the peptide backbone groups surrounding G302. Both the T304I and Q256L mutations can thus stabilize the conformation of the M^pro^ C-terminus that is required for its proper placement in the active site of another M^pro^ molecule for cleavage.

These interactions are also responsible for some active site conformational differences between the triple mutant and the WT. When comparing the triple mutant to a previously published M^pro^ C145A complex structure where the peptide substrate is also observed in the active site^17^, a distinct widening of the active site pocket is observed (Figure 2c). It appears that the Q256-containing helix nudges the C-terminal peptide towards the backbone of residues 166-168 in the active site of the adjacent protomer (that the P3-P5 substrate residues normally stack against) and that they consequently also flex outwards, with the most pronounced shifts occurring at F167 and P168. This outward shift of the backbone atoms of residues 166-168 affords extra room in the active site and allows the substrate binding orientation to shift slightly from the previous complex structure at the P2-P6 positions. The shift of the backbone also appears to place residue A166 close to the S1 residue of the other M^pro^ protomer in the same dimer, which would likely cause steric clashes between the WT E166 side chain and the S1 residue. This suggests that the E166A mutation can facilitate the conformational change observed in the triple mutant, as the smaller alanine side chain prevents such potential steric clashing. It may also explain the synergy observed between the L50F and the E166A/L167F mutations. Specifically, L50F improved the activity of the E166A/L167F mutant by 6-fold in the L50F/E166A/L167F triple mutant, in comparison to the 3-fold activity difference between the L50F single mutant and WT.

Aside from L50F, the triple mutant structure further illustrates the contribution of L167F to the binding of the nsp5-6 substrate. In this structure, V303 is sandwiched between F305 from the same M^pro^ molecule and F167 from the active site of the neighboring protein (Figure 2b). It would form significantly more interactions than those involving the equivalent T303 in the nsp4-5 substrate which also has L305 in the placement of F305. These interactions may be the reason for the difference in the L167F activity change vs. WT when using the two peptide substrates (Figure 1d).

### L50F single mutant crystal structure

In the 2.21 Å resolution crystal structure of M^pro^ L50F single mutant, the F50 side chain is also involved in extensive intermolecular hydrophobic interactions that help place the C-terminus of the adjacent monomer in the active site. However, some interesting differences exist compared with the triple mutant structure. First, when the two interacting dimers are constructed using crystallographic symmetry, there is a pseudo two-fold symmetry with each dimer placing the C-terminus of one protomer in the active site of a protomer from the other dimer (Figure 3a, Supplementary Data Figure 4). In both dimer-dimer interfaces of the L50F single mutant structure, F50 is involved in similar hydrophobic interactions with P252, F294, and V297, although additional contacts are made with P293 and I249 at the chain B dimer interface (Figure 3a). These hydrophobic interactions are different from those in the triple mutant structure involving V212, L220, T257, and I259. Second, the M^pro^ substrate C-terminus adopts a non-canonical conformation in the active site. Typically, the substrate enters from the bottom of the catalytic pocket with the substrate P3-P5 positions placed along the enzyme backbone from E166-P168 and the P4 side chain in the S4 pocket. However, in the L50F structure, the substrate enters from the side of the active site, with the substrate P3-P5 side chains “horizontal” to the enzyme active site (Figure 3b). In this new conformation, the P2 side chain (F305) is placed in the canonical S2 pocket. While Q306 resides near the S1 pocket, its position and conformation are slightly different from those in the triple mutant structure (Figure 3c), with the terminal carboxylate group placed outside the oxyanion hole. Thus, this conformation does not represent the product conformation immediately after peptide bond cleavage. Nevertheless, the P2 F305 and P1 Q306 adopt the superimposable configurations in both structures suggesting that P1-P2 substituents are more important than P3- P5 substituents in forming the ES complex. It is unclear whether substrate peptides entering the active site through this alternative conformation can be properly cleaved by the enzyme, although it is possible that C145 may not be able to access the scissile peptide bond if the alternative peptide conformation is maintained. We hypothesize that while the protein-protein interactions observed in the L50F single mutant structure can help place the C-terminal cleavage site in the enzyme catalytic center, this state does not represent the productive configuration required for the peptide bond cleavage, and additional conformational changes may be required for the reaction to proceed. In fact, in a previously published L50F structure (PDB 8DKZ)^12^ where a similar dimer of dimers configuration is observed and the substrate C-terminal peptide adopts the typical conformation with Q306 properly nestled in the S1 pocket, F50 only forms limited interactions with V212 of the adjacent substrate M^pro^. There are significantly fewer contacts than observed in our L50F single mutant or L50F/E166A/L167F triple mutant structures. It is possible that our L50F single mutant structure represents the initial encounter between the enzyme and substrate protein as facilitated by F50, and the previously published L50F structure (8DKZ) resembles the dimer of dimer configuration in the productive ES complex following some thermal motions after the initial encounter. It is consistent with the synergy between the L50F and E166A mutation and suggests that the optimal interactions involving the C-terminal peptide (as observed in the canonical binding mode) and F50 (as observed in our L50F or L50F/E166A/L167F structures) may not co-exist in the presence of the E166 side chain. It is also worth mentioning that similar dimer of dimers have also been observed in M^pro^ WT or C145A mutant crystal structures (PDB 7E5X, 7KHP)^18,19^ in addition to the aforementioned previously published L50F structure (8DKZ), where the C-terminus of one dimer is placed in the active site of the other dimer. The average surface area buried at the dimer interface is 1811 Å^2^ and 1375 Å^2^ for our L50F and L50F/E166A/L167F mutant structures respectively, in comparison to 902 Å^2^ (7E5X), 1090 Å^2^ (7KHP), and 866 Å^2^ (8DKZ) in these previously published structures, again highlighting the enhanced protein-protein interactions promoted by the L50F mutation revealed in our structures (Supplementary Data Figure 5).

**Figure 3.**
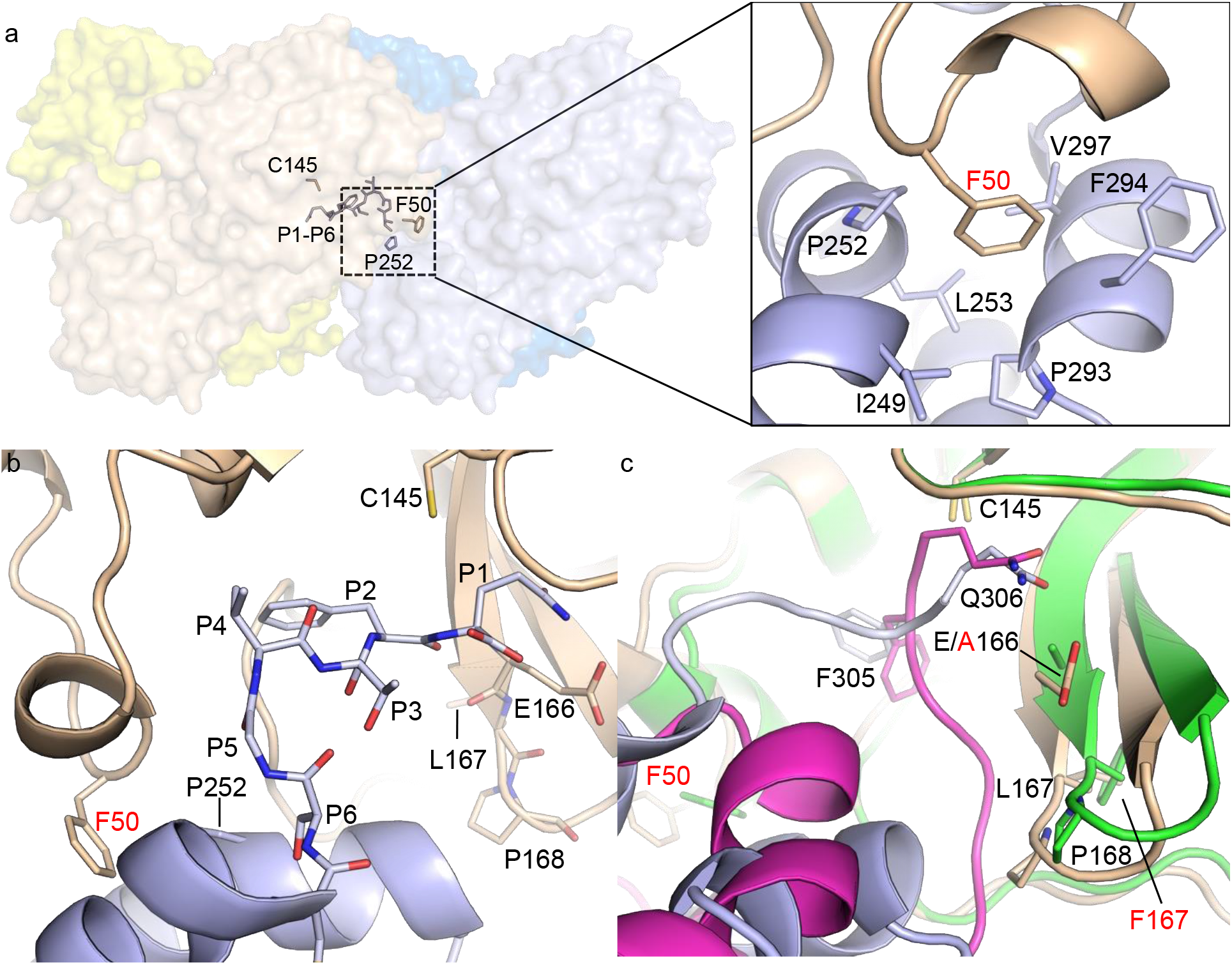
M^pro^ L50F single mutant. **a)** L50F single mutant dimer of a dimer (yellow/wheat, blue/light blue), showing P1-P6 (light blue) of the C-terminus bound in the chain B (wheat) active site. Zoomed in view shows the interactions of the enzyme F50 with the substrate hydrophobic pocket. A view of the chain A interactions can be found in Supplementary Data Fig. 4. Mutation is noted in red text. **b)** Binding of P1-P6 in the L50F chain B active site. The substrate is shown in light blue and the enzyme is shown in wheat. Mutation is noted in red text. **c)** Comparison of the binding modes of P1-P6 in the L50F single mutant (light blue/wheat) vs. the L50F/E166A/L167F triple mutant (magenta/green). Mutations are noted in red text.

In addition to L50F, the T21I and P252L mutations are two of the most frequently observed pathways leading to nirmatrelvir resistance in combination with active site mutations. The observation of P252 in the dimer-dimer interface of the L50F single mutant structure indicates that this residue may also affect protein-protein interactions in the ES complex. In the triple mutant structure, T21 also finds itself positioned on the edge of a hydrophobic pocket at the dimer-dimer interface, involving L67 from the same protomer and L232 and M235 from the other adjacent protomer (Supplementary Data Figure 6). The effect of the T21I mutation on the protein-protein interactions at this interface may be difficult to predict and possibly not significant due to its peripheral location. However, we hypothesize that T21I may affect protein-protein interactions in other ES complexes, including possibly involving other nsp6 residues during self-cleavage at the C-terminus or nsp4 residues at the N-terminus. In addition, other resistance mutations from viral passage assays are also found at the dimer-dimer interface of the triple mutant structure, such as A191V, A193P, Q256L, T304I, and P252L (Supplementary Data Figure 6)^15^. Together with the protein-protein interactions enhanced by L50F, these observations lend support to the likely biological relevance of this dimer-dimer interface and suggest the compatibility of these mutations with one another in conferring resistance.

As SARS-CoV-2 continues to evolve and mutate, it will be important to understand the mechanisms behind M^pro^ resistance to our current best treatments and identify the mutations that are critical to this resistance, as any variants that emerge with such mutations will be of particular concern to the global health community and necessitate careful monitoring. Our results demonstrate that distal protein-protein interactions, away from the active site, contribute significantly to M^pro^ substrate binding, and mutations at these locations can dramatically alter M^pro^ activity and the fitness of viral replication. This study provides a structural explanation for the role of L50F in promoting M^pro^ protein-protein interactions and restoring the reduced enzymatic activity caused by active site drug-resistant mutants such as E166A/L167F. Aside from M^pro^ self-cleavage, it is possible that mutations like L50F can also enhance the interactions between M^pro^ and other nsp proteins on the viral polyproteins. Our results, including those comparing nsp5-6 and nsp4-5 peptide substrates, highlight the need to investigate the effects of resistance mutations on specific enzyme-substrate interactions inside and outside the active site for different cleavage sites, rather than using one generic peptide substrate.

## Methods

### Fluorescence gel assay fusion protein construct and purification

The M^pro^ C145A (C145A/L50F) and *C. difficile* PBP3 42-554 fusion protein was inserted into the pETGSTSumo vector. The cleavage site, SAVKRT, was inserted between M^pro^ and PBP3 42-554. The expression constructs were transformed into Rosetta (DE3) pLysS cells. A single colony was picked and grew in LB (Luria-Bertani) media supplemented with 50 µg/mL kanamycin and 35 µg/mL chloramphenicol at 37 °C overnight. The overnight culture was then added into 1 L media at 1:100 and incubated at 37 °C until the OD_600_ reached 0.4. Protein expression was induced using 0.5 mM IPTG at 20 °C overnight. Cells were harvested by centrifugation at 5,000 x *g* for 10 min. The cell pellet was resuspended and disrupted by sonication in buffer A (20 mM Tris-HCl pH 8.0, 300 mM NaCl, 20 mM imidazole, 10 % glycerol), followed by ultracentrifugation at 45,000 x *g* for one hour. The supernatant was then loaded onto a HisTrap HP affinity column and eluted by linear gradient into buffer B (20 mM Tris-HCl pH 8.0, 300 mM NaCl, 500 mM imidazole, 10 % glycerol). The His tagged protein was pooled and buffer exchanged into cleavage buffer (20 mM Tris-HCl pH 8.0, 100 mM NaCl, and 10 % glycerol). The Sumo tag was removed by ULP1 incubation at 4 °C overnight. The sample was then loaded onto a HisTrap HP column for cleanup. The flowthrough was further purified using a HiPrep 16/60 Sepharcyl S-300 HR size exclusion column in storage buffer (20 mM Tris-HCl pH 8.0, 200 mM NaCl, 1 mM DTT).

### Fluorescence gel assay

Purified untagged M^pro^ C145A-PBP3 42-554 was diluted in assay buffer (20 mM Tris-HCl pH 8.0, 200 mM NaCl) and labeled with 1.5-fold Bocillin at room temperature for 30 minutes. 10 µM labeled M^pro^ C145A-PBP3 42-554 was incubated with 10 µM of different M^pro^ mutants for the indicated times. The reaction was stopped by using 2x SDS-PAGE loading buffer without dye. The samples were then loaded onto Tris-Glycine 7.5% SDS-PAGE gels. The intensity of the protein bands was analyzed using a ChemiDoc XRS+ and the ImageJ software. The initial rate was analyzed by SigmaPlot and reported as 10^-7^ M/min. Rates are given as mean±s.d. for two biologically independent replicates.

### M^pro^ FRET enzymatic assay

The M^pro^ nsp4-5 and nsp5-6 FRET substrates were synthesized using the Fmoc solid peptide synthesis^20^. The sequences are: nsp4-5/Dabcyl-K-TSAVLQ↓SGFRKM-E(Edans)-CONH_2_; nsp5-6/Dabcyl-K-SGVTFQ↓SAVKRT-E(Edans)-CONH_2_.

K_m_ and V_max_ of SARS-CoV-2 M^pro^ mutants were performed using the optimized concentration (the concentration that allows the initial velocity to saturate in the testing substrate concentration range). The final concentrations of substrate ranges from 0.78-200 µM. The reaction was carried out in a total volume of 100 µL reaction buffer containing 20 mM HEPES pH 6.5, 120 mM NaCl, 0.4 mM EDTA, 4 mM DTT, and 20 % glycerol. The signal was detected using a BioTek Cytation 5 imaging reader (Agilent) with the excitation at 360/40 nm and emission at 460/40 nm. The reaction was monitored every 70 seconds. The initial velocity of proteolytic activity was determined by linear regression of the first 600-1000 seconds of the kinetic progress curves. The K_m_ and V_max_ were calculated by plotting the initial velocity against FRET substrate concentrations using the classic Michaelis-Menten equation (*Y = V_max_*X/(K_m_ + X)*, *X* = substrate concentration; *Y* = enzyme velocity) in the Prism 8 software.

For K_i_ determination, the optimized mutant SARS-CoV-2 protein concentration (the concentration that gives at least 1-hour linear initial velocity) was mixed with 20 µM FRET substrate and various concentrations of nirmatrelvir in 100 µL volume of reaction buffer. The reactions were monitored every 70 seconds for 3 hours. The initial velocity was determined for the first hour by linear regression. The K_i_ was determined by plotting the initial velocity against inhibitor concentrations using the Morrison equation for tight binding *(Y =V_0_*(1 − ((((E_t_ + X + (K_i_*(1 + (S/K_m_)))) − (((E_t_ + X +(K_i_*(1 + (S/K_m_))))^2^^) −4*E_t_*X)*^0*.5^*^ *))/(2*E_t_)))*, *X* = inhibitor concentration; *Y* = enzyme velocity; *E_t_* = enzyme concentration; *V_0_* = enzyme velocity in the absence of inhibitor). The reported values were the average of two replicates ± standard error with a 95% confidence interval calculated as SE = (upper limit − lower limit)/3.92.

### M^pro^ mutagenesis, protein expression, and purification

M^pro^ mutants were generated using the QuikChange® II Site-Directed Mutagenesis Kit from Agilent (Catalog #200524), using pET-SUMO-M^pro^ (from strain BetaCoV/Wuhan/WIV04/2019) plasmid as the template.

M^pro^ mutant proteins were expressed and purified as previously described, with minor modifications. Plasmids were transformed into RosettaPlyss (DE3) competent cells, and bacterial cultures overexpressing the target proteins were grown in LB media containing 50 µg/mL of kanamycin and 35 µg/mL chloramphenicol at 37 °C. Expression of the target protein was induced at an OD_600_ of 0.6-0.8 by the addition of isopropyl β-d-1-thiogalactopyranoside (IPTG) to a final concentration of 0.5 mM. The cell culture was incubated at 20 °C for 12-16 hrs. Bacterial cultures were harvested by centrifugation (5,000 × *g*, 10 min, 4 °C) and resuspended in His buffer A (20 mM Tris pH 8.0, 300 mM NaCl, 40 mM imidazole, 10 % glycerol). Bacterial cells were lysed by alternating sonication (10% amplitude, 10 s on/15 s off). The lysed cell suspension was clarified by centrifugation (45,000 × *g*, 60 min, 4 °C) and the supernatant loaded onto a HisTrap HP column. The column was thoroughly washed with 60 mM imidazole in lysis buffer. The protein was eluted by linear gradient, 1-100 %, using His buffer B (20 mM Tris pH 8.0, 300 mM NaCl, 500 mM imidazole, 10 % glycerol). The eluted protein was pooled and buffer exchanged into cleavage buffer (20 mM Tris pH 8.0, 100 mM NaCl, 10 % glycerol), and then incubated with ULP1 protease at 4 °C overnight. The sample was then loaded onto a HisTrap HP column, and the flowthrough was collected and concentrated. The flowthrough was further purified using a HiPrep 16/60 Sepharcyl S-300 HR size exclusion column in storage buffer (20 mM Tris-HCl pH 8.0, 200 mM NaCl, 1 mM DTT).

### M^pro^ crystallization and structure determination

The M^pro^ mutants were crystallized as previously described^7^. Briefly, M^pro^ mutants were diluted to 5 mg/mL in storage buffer. Crystals were grown by hanging drop in 25 % PEG 3350, 0.1 M Na/K tartrate, 0.005 M MgCl_2_, by mixing 1.5 µL of the protein solution with 1.5 µL of the crystallization condition. Crystals grew after 1-3 days of incubation at 20 °C. Crystals were transferred into a cryoprotectant solution (crystallization condition supplemented with 20 % glycerol) and flash-frozen in liquid nitrogen.

X-ray diffraction data were collected at cryogenic temperature (100 K) at the Structural Biology Center (SBC) 19-ID beamline at the Advanced Photon Source (APS) in Argonne, IL, using a Pilatus3 6M detector and a wavelength of 0.97918 Å. Data were processed with HKL-3000 and CCP4, and PHASER was used for molecular replacement using a previously solved SARS-COV-2 M^pro^ structure (PDB 6WTT) as the reference model. Model building and structure refinement was completed using the CCP4 suite^21^, Coot^22^, and the PDB REDO server (pdb-redo.edu)^23^. All images were generated using the PyMOL Molecular Graphics System (Schrödinger, LLC).

Full crystallographic statistics are provided in Supplementary Data Table 1.

### Buried surface area calculation

The total solvent accessible surface area (SASA) was computed for isolated dimers and the specific SASA restricted against the adjacent dimer. The dimer-dimer buried surface area (ddBSA) was then calculated for each dimer as the difference between these two values, and then averaged. The VMD program’s implementation of the maximal speed molecular surfaces algorithm (measure SASA command, restrict option)^24,25^ with a probe radius of 1.4 Å was used for this analysis. All non-protein moieties were stripped from the coordinate files. Dimer of dimer complexes were generated using symmetry operations in PyMOL if not part of the asymmetric unit.

## Data Availability

The X-ray crystal structures used in this study have been deposited in the Protein Data Bank (PDB) with accession codes 8U4Y (M^pro^ L50F) and 8U25 (M^pro^ L50F/E166A/L167F).

## Acknowledgements

This research was supported by the National Institutes of Health (NIH) grant AI158775 to J. W. and Y. C. Results shown in this report are derived from work performed at Structural Biology Center funded by the U.S. Department of Energy (DOE), Office of Biological and Environmental Research, and operated for the DOE Office of Science at the Advanced Photon Source by Argonne National Laboratory under Contract No. DE-AC02-06CH11357. We thank the scientists and staff at SBC for their assistance with X-ray diffraction data collection.

## Author Contributions

E.M.L, J.W, and Y.C wrote the manuscript; X.Z, A.F, and L.M.C.J carried out the gel-assay; H.T. performed the FRET enzymatic assay; E.M.L and A.J.B crystallized proteins; E.M.L and N.Ksolved and deposited protein structures; J.J.M carried out computational analysis of the crystal structures.

## Additional Information

Correspondence should be addressed to Yu Chen at.

