## Supplementary Material for "Distal Protein-Protein Interactions Contribute to SARS-CoV-2 Main Protease Substrate Binding and Nirmatrelvir Resistance"

**This PDF file includes:**

Supplementary Data Figures 1-6

Supplementary Data Table 1

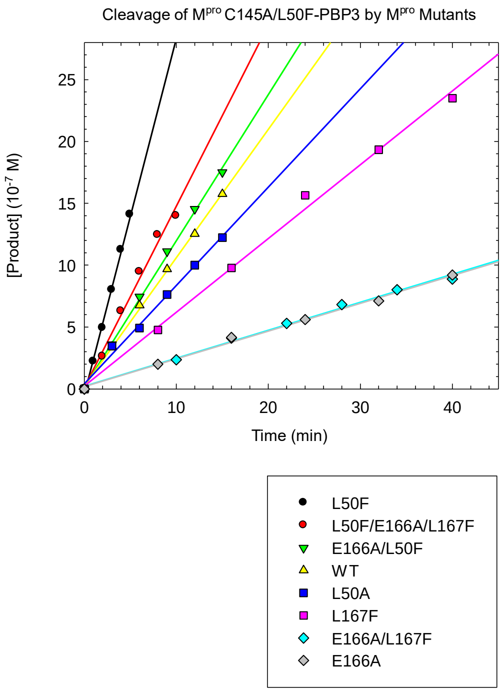


**Supplementary Data Figure 1.** Activity of C145A/L50F-PBP3 against multiple $M^{pro}$ mutants.

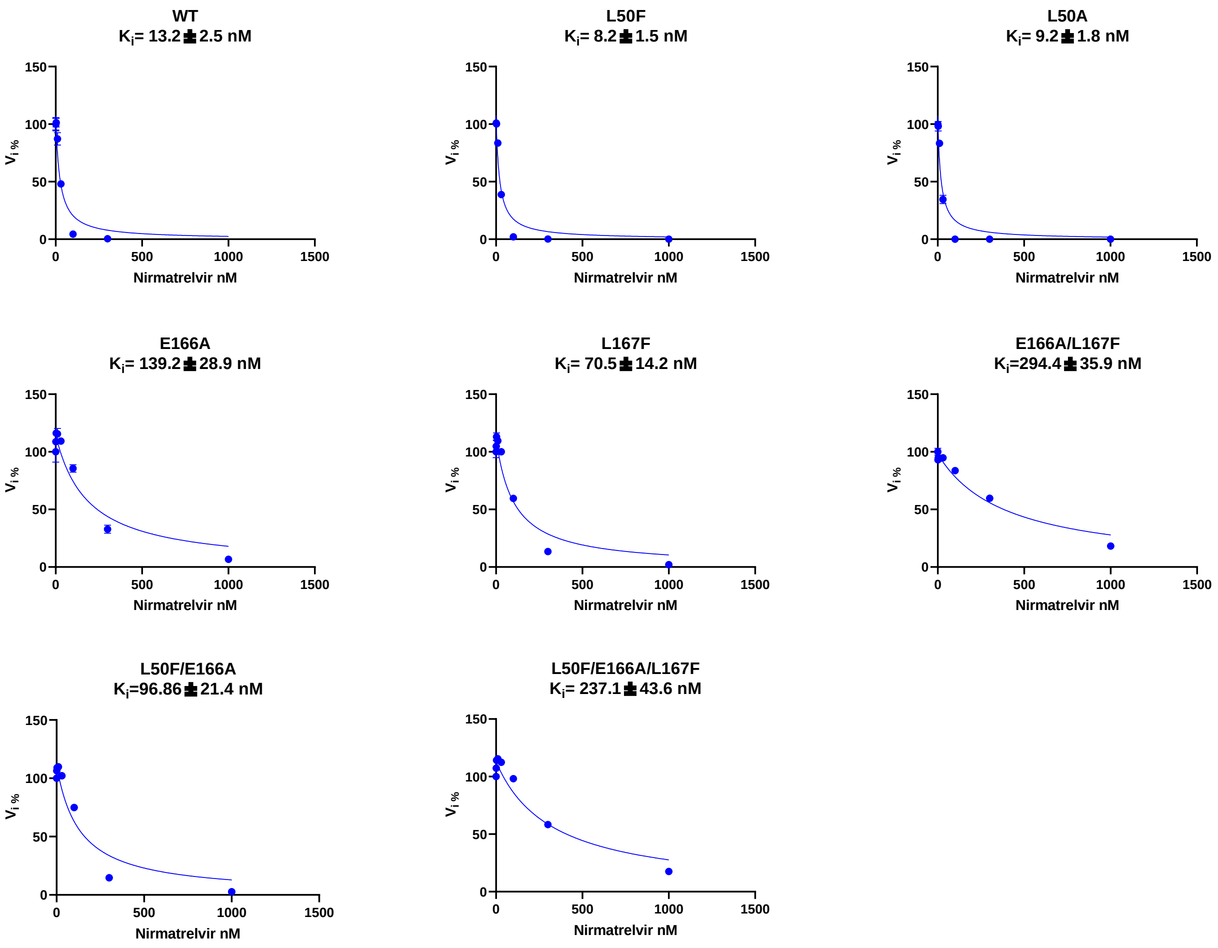


**Supplementary Data Figure 2.** Dose response curves of nirmatrelvir inhibition vs. nsp5/6.

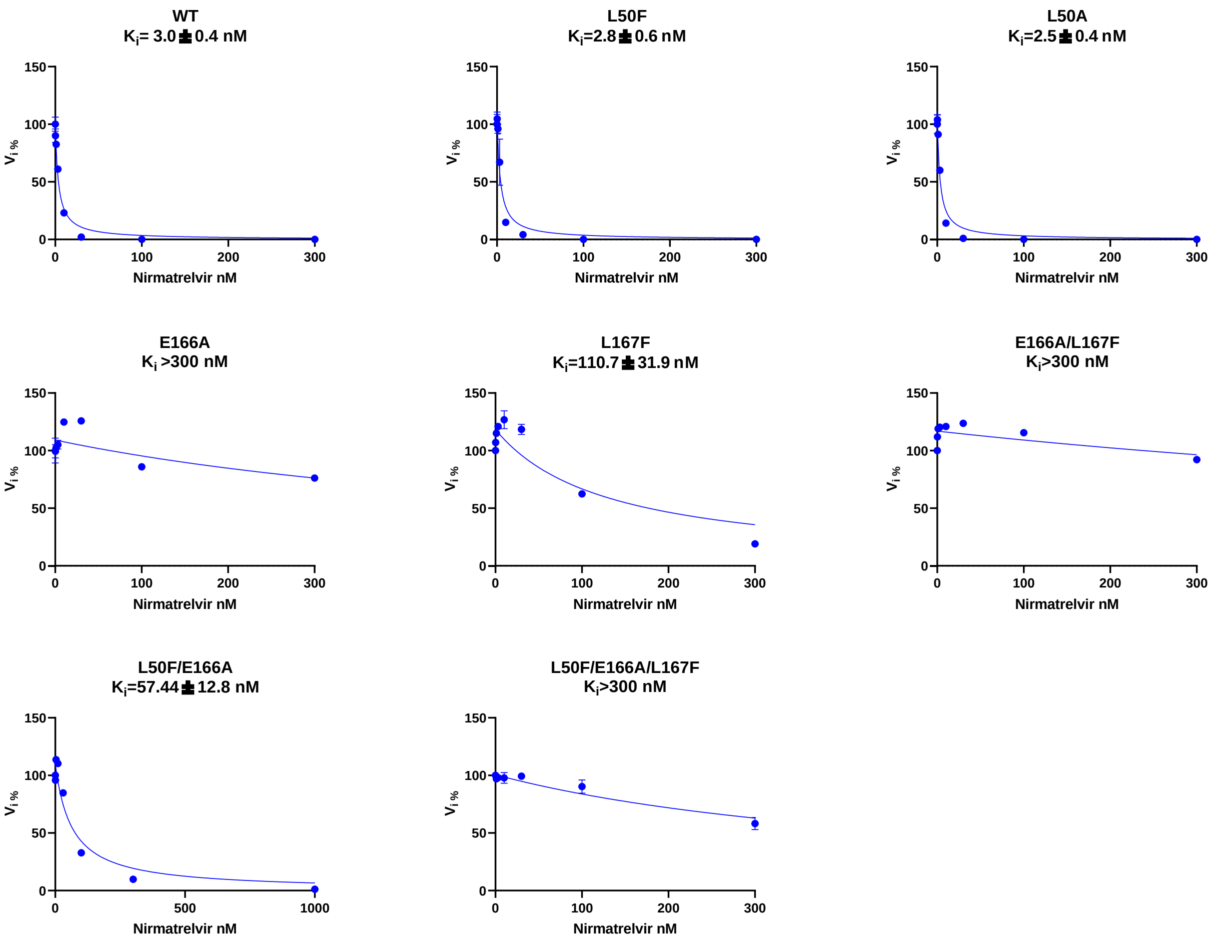


**Supplementary Data Figure 3.** Dose response curves of nirmatrelvir inhibition vs. nsp4/5.

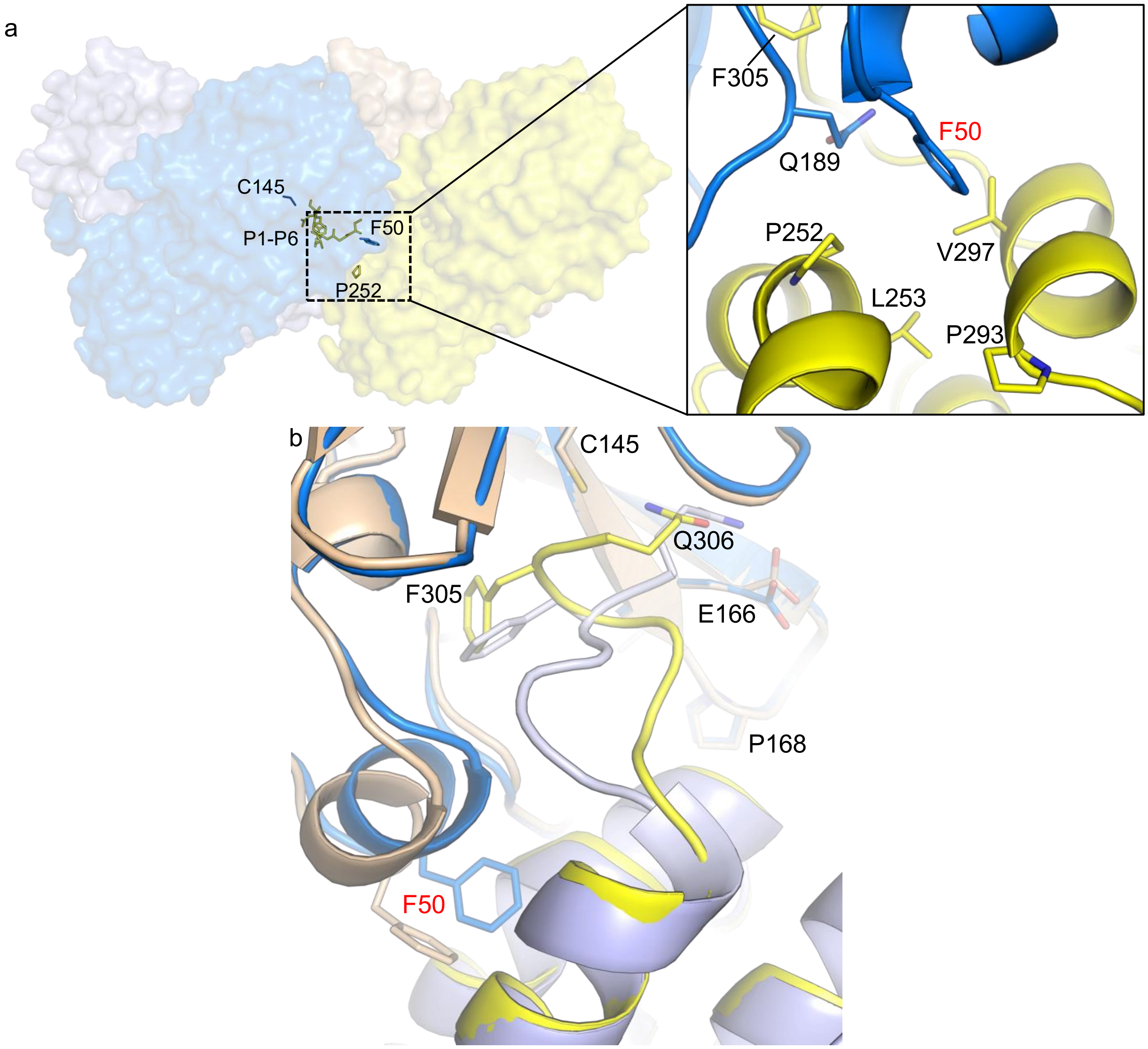


**Supplementary Data Figure 4. a)** L50F single mutant dimer of a dimer (blue/light blue, yellow/wheat), showing P1-P6 (yellow) of the C-terminus bound in the chain A (blue) active site. Zoomed in view shows the interactions of the enzyme F50 with the substrate hydrophobic pocket. Mutation is noted in red text. **b)** Comparison of the binding pose of P1-P6 in the active site of chain A (blue/yellow) vs. chain B (light blue/wheat). Mutation is noted in red text.

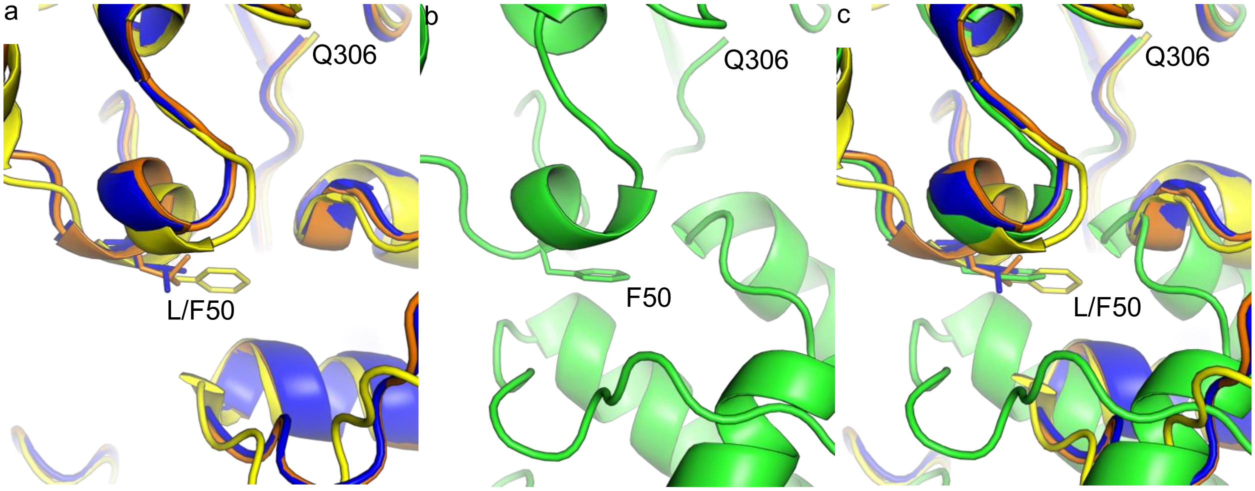


**Supplementary Data Figure 5.** Enhanced F50 interactions in the $M^{pro}$ L50F/E166A/L167F triple mutant. **a)** Aligned structures of $M^{pro}$ in complex with its C-terminal cleavage sequence (orange, WT, PDB 7E5X; blue, WT, PDB 7KHP; yellow, L50F, PDB 8DZK). **b)** $M^{pro}$ L50F/E166A/L167F triple mutant in complex with its C-terminal cleavage sequence (green, PDB 8U25). **c)** Alignment of the triple mutant (green) with the $M^{pro}$ structures discussed in panel a (orange, blue, yellow).

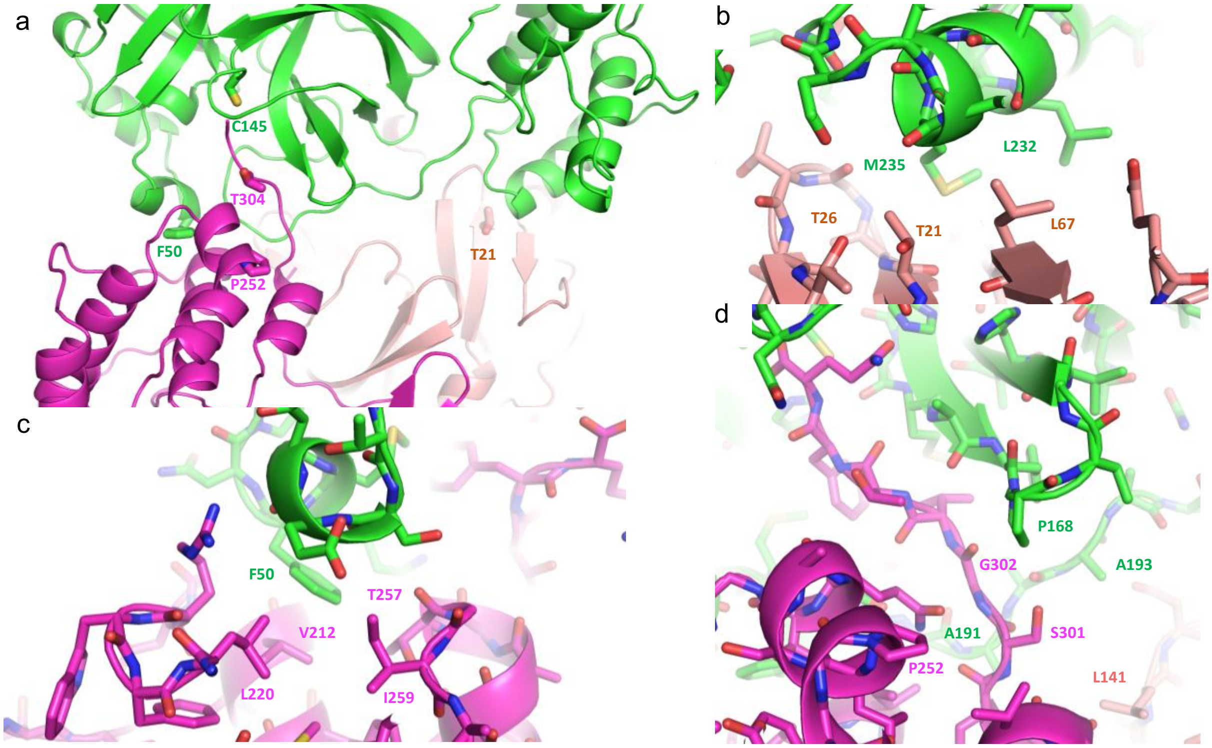


**Supplementary Data Figure 6.** Resistance mutations at the dimer-dimer interface. **a)** The dimer-dimer interface of the L50F/E166A/L167F triple mutant, involving one protomer (green) from the enzyme dimer and two protomers (magenta and salmon) from the substrate dimer. The side chains of the catalytic C145 and F50 from the enzyme dimer, and three resistance mutation hot spots from the substrate dimer, are shown in stick. Most of the main non-active site resistance mutation hot spots from the viral passage assays (e.g., T21, P252) are found at this interface. **b)** Protein-protein interface near T21. **c)** Protein-protein interface near F50. **d)** A potential resistance hot spot near P168. A191V, A193P, and Q256L were also identified in the viral passage assay. P168S is the most abundant natural variant at residue 168.

**Supplementary Data Table 1. X-ray Data Collection and Refinement Statistics**

| | L50F[a] | L50F/E166A/L167F[a] |
|---|---|---|
| | PDB Code: 8U4Y | PDB Code: 8U25 |
| **Data collection** | | |
| Space group | $P2_1$ | $P2_1$ |
| Cell dimensions | | |
| $a$, $b$, $c$ (Å) | 48.23, 105.47, 53.79 | 71.12, 87.23, 107.56 |
| $\alpha$, $\beta$, $\gamma$ (°) | 90, 104.50, 90 | 90, 104.49, 90 |
| Resolution (Å) | 50-2.22 (2.26-2.21)[b] | 50-2.23 (2.27-2.23) |
| No. of Reflections | 25847 (1265) | 60743 (2876) |
| $R_{merge}$ (%) | 13.3 (51.7) | 14.7 (51.8) |
| $I$ / $\sigma I$ | 17.3 (2.41) | 10.2 (2.14) |
| Completeness (%) | 99.4 (97.7) | 97.9 (94.6) |
| Redundancy | 6.2 (4.4) | 3.9 (3.4) |
| **Refinement** | | |
| Resolution (Å) | 46.74-2.22 | 47.71-2.23 |
| No. reflections | 24493 | 57640 |
| $R_{work}$ / $R_{free}$ | 17.9/23.9 | 18.0/23.1 |
| No. atoms | | |
| Protein | 4748 | 9500 |
| Ligand/ion | 0 | 0 |
| Water | 82 | 506 |
| *B*-factors | | |
| Protein | 47.2 | 32.09 |
| Ligand/ion | 0 | 0 |
| Water | 41.75 | 31.27 |
| R.m.s. deviations | | |
| Bond lengths (Å) | 0.005 | 0.006 |
| Ramachandran Plot | | |
| Favored Region (%) | 98.8 | 98.9 |
| Allowed Region (%) | 0.9 | 0.8 |
| Outlier Region (%) | 0.2 | 0.4 |

[a] Single crystal used for dataset.

[b] Values in parentheses are for highest-resolution shell.
